# Neuron-specific cilia loss alters locomotor responses to amphetamine

**DOI:** 10.1101/2020.03.16.994087

**Authors:** Carlos Ramos, Jonté B. Roberts, Kalene R. Jasso, Tyler W. Ten Eyck, Barry Setlow, Jeremy C. McIntyre

**Author notes:** Authors contributed equally (Carlos Ramos, Jonté B. Roberts). Corresponding Author: Jeremy C. McIntyre, PO Box 100244, Gainesville, FL 32610.

## Abstract

The neural mechanisms that underlie responses to drugs of abuse are complex, and impacted by a number of neuromodulatory peptides. Within the past ten years it has been discovered that several of the receptors for neuromodulators are enriched in the primary cilia of neurons. Primary cilia are microtubule-based organelles that project from the surface of nearly all mammalian cells, including neurons. Despite what we know about cilia, our understanding of how cilia regulate neuronal function and behavior is still limited. The primary objective of this study was to investigate the contributions of primary cilia on specific neuronal populations to behavioral responses to amphetamine. To test the consequences of cilia loss on amphetamine-induced locomotor activity we selectively ablated cilia from dopaminergic or GAD2-GABAergic neurons in mice. Cilia loss had no effect on baseline locomotion in either mouse strain. Both female and male mice lacking cilia on dopaminergic neurons showed significantly reduced responses to acute administration of 3.0 mg/kg amphetamine compared to wildtype mice. In contrast, changes in the locomotor response to amphetamine in mice lacking cilia on GAD2-GABAergic neurons were primarily driven by reductions in locomotor activity in males. Following repeated amphetamine administration (1.0 mg/kg/day over 5 days), mice lacking cilia on GAD2-GABAergic neurons exhibited enhanced sensitization of the locomotor stimulant response to the drug, whereas mice lacking cilia on dopaminergic neurons did not differ from their wildtype controls. These results indicate that cilia play neuron-specific roles in both acute and neuroplastic responses to psychostimulant drugs of abuse.

## 1. Introduction

The primary cilium is a microtubule-based organelle that projects from the surface of nearly all mammalian cell types including neurons [1-3].Cilia have emerged over the past 15 years as important regulators of neuronal function. All neurons have cilia, and diseases caused by mutations in cilia-specific genes, known collectively as ciliopathies, are accompanied by deficits in cognition and motivated behavior [4-15]. Across organ systems, cilia play a variety of signaling roles, largely related to sensing extracellular cues. Consistent with these roles, neuronal cilia express a variety of G protein-coupled receptors (GPCRs), several of which are rarely expressed outside of cilia [16-18]. Importantly, several of these GPCRs modulate behaviors related to drugs of abuse. For example, the receptor for melanin-concentrating hormone (MCHR1) modulates responses to psychostimulants, and the orphan GPCR, GPR88, modulates alcohol intake [16, 19-22]. To date, however, the role or necessity of neuronal cilia in responses to drugs of abuse is essentially unknown.

Nearly all drugs of abuse target (either directly or indirectly) dopaminergic neurons in the midbrain and their projections to the nucleus accumbens (NAc), which is largely comprised of GABAergic neurons. As a first step toward determining how neuronal cilia impact responses to drugs of abuse, we evaluated locomotor behavior following both acute and repeated administration of amphetamine, the prototypical drug of abuse, in mice engineered to lack neuronal cilia on either dopaminergic or GAD2-GABAergic neurons. Locomotor response to amphetamine likely does not directly reflect rewarding or reinforcing properties of the drug; however, such measures can provide evidence for drug responsivity and drug-induced neuroplasticity that may be relevant for some aspects of substance use [23, 24]. In addition, evaluation of cilia on multiple neuron types can help localize specific cilia populations most relevant for drug responsivity. Notably, cilia in midbrain dopaminergic and NAc GABAergic neurons express different complements of neuromodulatory receptors. For example, both MCHR1 and GPR88 are enriched in cilia in NAc, but are not detected in cilia on midbrain dopaminergic neurons [16, 17, 25]

The results show that cilia ablation on either GAD2-GABAergic or dopaminergic neurons attenuates the locomotor stimulant effect of acute amphetamine. In contrast, cilia ablation on GAD2-GABAergic neurons enhances sensitization of the locomotor stimulant effect of repeated amphetamine, whereas ablation on dopaminergic neurons has no effect. Taken together, these findings indicate cell-specific roles for cilia in regulating neuronal responses to amphetamine, and support further investigation of interactions between cilia and drugs of abuse, as well as the contributions of this organelle to substance use.

## 2. Materials and methods

### 2.1 Animals

Neuron-specific knockouts were generated through conditional deletion of the cilia gene *Ift88* (intraflagellar transport protein 88) [26, 27]. Mice with floxed alleles of *Ift88* (Ift88^tm1Bky^) were crossed to strains expressing cell-specific Cre Recombinase (Cre); Gad2^IresCre^ (Jax stock 010802) mice for GAD2-GABAergic neurons (referred to as GAD2:Cilia mice), and DAT^iresCre^ mice (Jax stock 006660) for dopaminergic neurons (referred to as DAT:Cilia mice) [26, 28, 29]. Breeding strategies and offspring genotypes are shown in Table 1. Mice were genotyped by extracting DNA from tail clippings with Extracta DNA Prep for PCR – Tissue (Quanta Biosciences) and specified products amplified using either GoTaq Green Mastermix (Promega) or 2x KAPA buffer. Mice of both sexes between 10 and 12 weeks of age were used for all experiments and group housed 3-5 per cage from weaning until use. Not all animals that were monitored for body weight were used for behavior experiments in this manuscript. The estrous cycle of female mice was not monitored for these experiments. Mice were maintained on a 12/12 light/dark cycle (lights on at 0700) with *ad lib* access to food and water. Experiments were performed between 0800 and 1700. All procedures were approved by the University of Florida Institutional Animal Care and Use Committee and followed NIH guidelines.

**Table 1.**
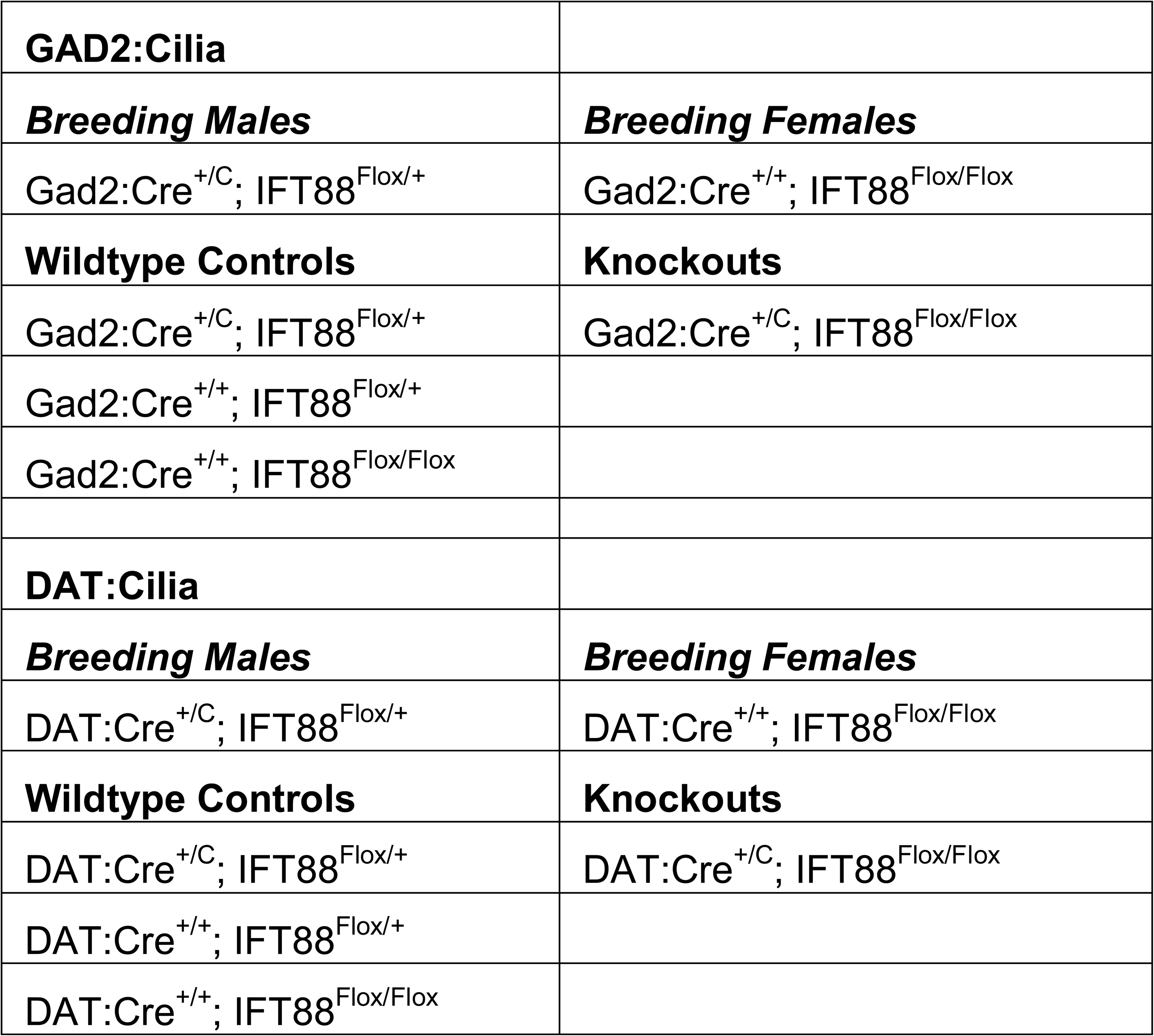
Genotypes of breeding animals and experimental animals used in this study. Males and females of both strains were used for all experiments.

### 2.2 Equipment and Behavior experiments

Locomotor activity was assessed in Med Associates activity monitoring chambers equipped with infrared beams to detect movement. The chambers were housed within sound-attenuating cubicles, and locomotor activity was assessed in the dark (lights off in the cubicles). To assess acute locomotor effects of amphetamine, mice were initially monitored for 1 h in the chambers, followed by i.p. injection of amphetamine (3 mg/kg) and monitoring for an additional hour. Locomotion and stereotypy were determined using Med Associates software and calculated in 5 min bins. The two primary measures of interest were distance traveled (calculated from the Euclidean distance of all ambulatory episodes and recorded in centimeters) and stereotypy (calculated as repeated breaks of the same infra-red beam). To assess locomotor sensitization to amphetamine, mice were initially monitored for 1 h in the chambers, followed by i.p. injection of amphetamine (1 mg/kg) and monitored for an additional hour on the first day. On days 2-5, the pre-injection monitoring period was omitted and mice were injected and placed in the chambers for 1 h. The amphetamine doses were chosen on the basis of data showing that they bracket the range in which both acute locomotor stimulation and locomotor sensitization are observed (e.g.,[30]), to allow detection of either increases and decreases in amphetamine-induced locomotion caused by cilia deletion.

### 2.3 Drugs

D-amphetamine sulfate was obtained from the NIDA Drug Supply Program. A 0.9% saline solution was used as vehicle. All doses were administered in a volume of 10.0□ml/kg body weight. Drug solutions were freshly prepared on each day of the experiments.

### 2.4 Immunohistochemistry

Mice were deeply anesthetized prior to cardiac perfusion with 4% paraformaldehyde (PFA). Dissected brains were incubated in 4% PFA overnight at 4°C then cryoprotected in 10%, 20% 30% sucrose for 1 hour, 1 hour, and overnight, respectively, at 4°C, and embedded in OCT compound (Tissuetek). Coronal sections of embedded tissues were cut at a thickness of 10-12 µm and mounted onto Superfrost Plus slides (Fisher Scientific). For immunostaining, sections were permeabilized and blocked with 0.3% Triton X-100 and 2% goat serum in PBS for 30 min. Primary antibodies were diluted in blocking buffer and applied to samples overnight at 4°C. Primary antibodies and concentrations used are shown in Table 2. For specific antibodies, MCHR1, an antigen retrieval step was included in which samples were steamed at 95°C in 6.5mM citric acid (pH 6.0) for 30min prior to blocking steps. Fluorescent-conjugated secondary antibodies (Alexa Probes, Invitrogen) were applied (at 1:1000 dilution) for 1 hour, followed by three washes with PBS. 4′,6-diamidino-2-phenylindole (DAPI) was applied for 5 min to stain nuclei, and samples were mounted using Prolong Gold (Invitrogen). Fixed tissue imaging was performed on a Nikon TiE-PFS-A1R confocal microscope equipped with a 488 nm laser diode with a 510-560 nm band pass filter, and 561 laser with a 575-625 nm band pass filter. A CFI Apo Lambda S 60 x 1.4 N.A. objective was used. Confocal *Z*-stacks were processed using NIH ImageJ software.

**Table 2.** Antibodies used for immunofluorescence.

| Target | Species | Company | Dilution |
| --- | --- | --- | --- |
| Anti-ADCY3 | Rabbit | EnCor, RPCA-ACIII | 1:2000 |
| Anti-ADCY3 | Mouse | EnCor, MCA-5J11 | 1:1000 |
| Ani-NeuN | Mouse | Millipore, AB5390 | 1:1000 |
| Anti-TH | Rabbit | Millipore, AB152 | 1:500 |
| Anti-GFP | Chick | Invitrogen, A10262 | 1:1000 |
| Anti-TH | Mouse | Invitrogen, A711649 | 1:200 |
| Anti-MCHR1 | Rabbit | Invitrogen, A711649 | 1:500 |

### 2.5 Statistics

Statistical analyses were conducted in SPSS 26.0. The effects of acute amphetamine were assessed separately in each strain of mice via multi-factor repeated measures ANOVA, with time bin as a within-subjects factor, and sex and genotype (Cilia^-ve^ vs. wild type) as between-subjects factors. Effects of sex and genotype on baseline locomotion (prior to amphetamine injections) were assessed in a similar manner. The effects of repeated amphetamine were also assessed separately in each strain using repeated measures ANOVA, with day as a within-subjects factor, and sex and genotype as between-subjects factors. For all analyses, *p*-values less than or equal to 0.05 were considered statistically significant.

## 3. Results

### 3.1 Validation of Cilia Loss

A single primary cilium projects from the cell body of nearly all neurons in the brain [31]. These structures are enriched with a number of g-protein coupled receptors (GPCRs) and other signaling molecules such as adenylyl cyclase 3 (ADCY3), which is a robust marker for neuronal cilia (Figure 1A). One of these GPCRs, MCHR1, shows differential expression in the NAc and VTA (Figure 1B), providing evidence that these structures may contribute to behaviors differently in specific brain regions. To assess the role of neuronal cilia in locomotor responses to amphetamine, we selectively ablated cilia from either GAD2-GABAergic or dopaminergic (DAT) neurons. In both mouse lines for this study, Cre-mediated *Ift88* excision occurs after neuronal differentiation. Removal of cilia from other neuronal strains, or in adult animals, has not produced noticeable changes in neuron survival up until at least 6 months of age [7, 27, 32, 33]. We confirmed cilia loss in both lines through immunohistochemistry for ADCY3. In both the GAD2:Cilia^-ve^ and DAT:Cilia^-ve^ mice, cilia are abolished from their respective cell types (Figure 2). Importantly, we did not notice overt deficiencies in survival of Cilia^-ve^ mice from either line, nor any evidence of obesity, which is a hallmark of many ciliopathies [4, 34, 35]. Instead, GAD2:Cilia^-ve^ mice showed a significant reduction in body weight compared to wildtype littermates at 10-12 weeks of age (main effect of genotype, F_(1,111)_=85.35, p<.001) that did not differ by sex (genotype x sex interaction, F_(1,111)_=2.20, p=.14, Figure 3A), whereas the DAT:Cilia^- ve^ mice did not differ from their wild type controls (main effect of genotype, F_(1,102)_=0.08, p=.78; genotype x sex interaction, F_(1,102)_=1.55, p=.22, Figure 3B).

**Figure 1.**
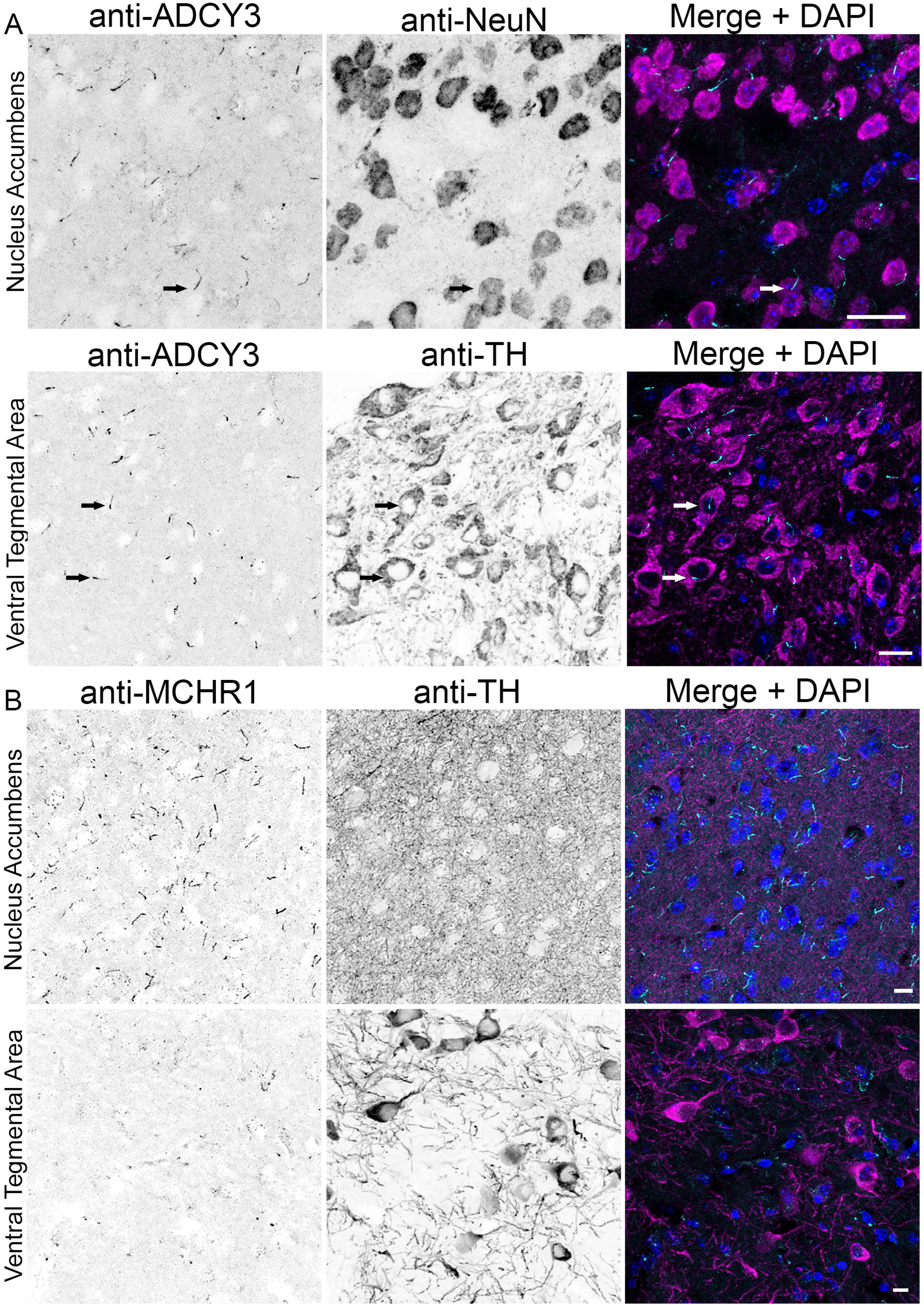
Identification of Cilia on primary neurons. (A) Immunofluorescence staining for ADCY3 in the nucleus accumbens (NAc, Top) and ventral tegmental area (VTA, bottom), shows cilia on neurons in both regions. Neurons are labelled with NeuN in the NAc and TH in the VTA. (B) Staining for the GPCR MCHR1, along with TH, in both regions shows a difference in the expression of MCHR1. Whereas abundant MCHR1 is detected in the NAc (top) sparse MCHR1 labeling is seen in VTA. Scale bars = 10μm

**Figure 2.**
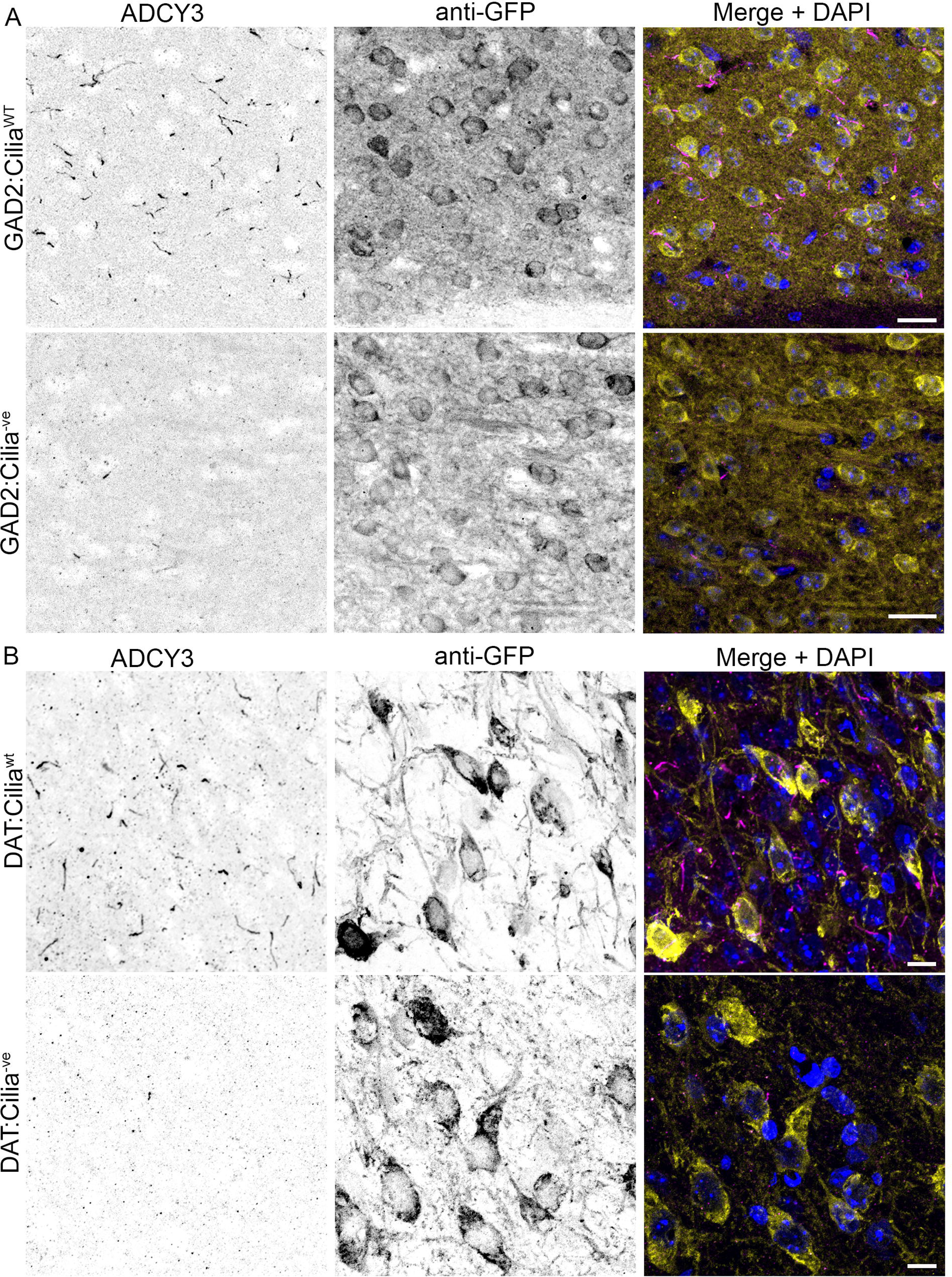
Cell specific expression of Cre results in cilia loss in Ift88^F/F^ mice. Immunofluorescence staining for the cilia marker, ADCY3, in WT and Cilia^-ve^ mice. To aid in identification of cell types, mice were produced that expressed a Cre dependent GCaMP6 from the Rosa26 locus. Immunostaining for GFP reveals GCaMP expression in both lines confirming Cre expression. (A) GAD2:Cilia^-ve^ mice (bottom) show GFP^+ve^ cells that lack ADCY3^+ve^ cilia. (B) Immunostaining showing the presence of cilia on DAT:Cre^+ve^ cells in WT mice (top), and the loss of ADCY3 staining in DAT:Cilia^-ve^ mice (bottom). Scale bars = 10μm

**Figure 3.**
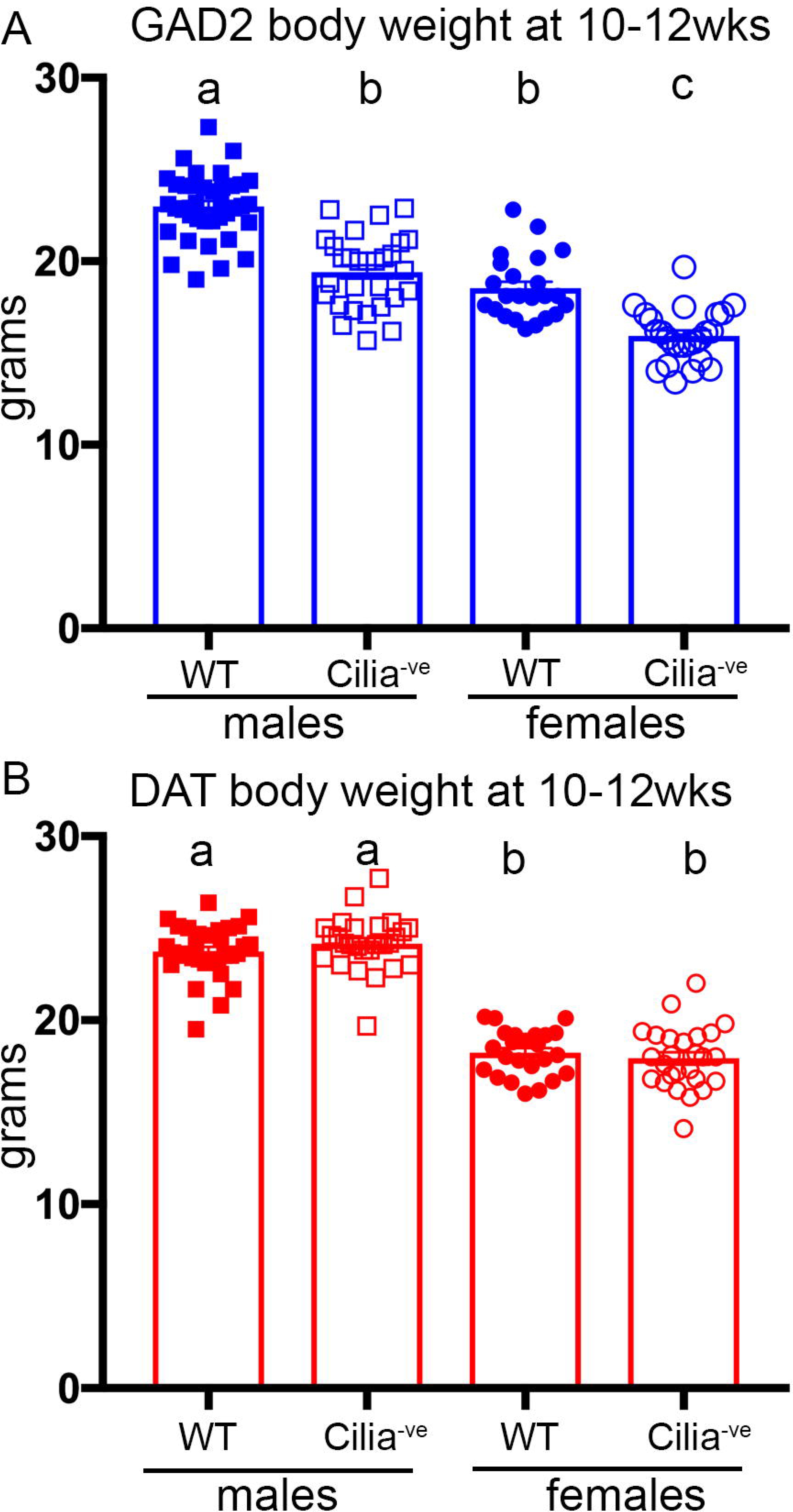
Mice lacking cilia on GAD2-GABAergic neurons weigh significantly less. Bar graphs showing average and individual weights of wildtype (WT) and Cilia^-ve^ male and female mice between 10 and 12 weeks of age. (A) Both male and female GAD2:Cilia^-ve^ mice weigh significantly less that wildtype littermates. (B) No differences in body weight are present in DAT:Cilia^-ve^ compared to wildtype mice. Columns with different letters are significantly different from each other.

### 3.2 Effects of genotype on baseline activity

To determine whether cilia loss on GAD2-expressing cells affects baseline activity, data from the 60 min prior to acute (3.0 mg/kg) amphetamine administration were evaluated. A 3-factor ANOVA (sex x genotype x time bin) conducted on the distance traveled measure revealed a significant effect of time bin (F_(11,418)_=95.63, p<.001), such that all mice decreased their distance traveled across bins, but no main effects or interactions involving genotype or sex (Fs<1.87, ps>.12)(Fig 4A, B). The same analysis conducted on the stereotypy counts also revealed a main effect of time bin (F_(11,418)_=66.04, p<.001), as well as a time bin x sex interaction such that the decrease in stereotypy counts across the session differed in males and females (F_(11,418)_=1.91, p=.04), but no main effects or interactions involving genotype (Fs<1.12, ps>.34)(Fig 4C, D). To determine whether cilia loss on DAT-expressing cells affects baseline activity, the same analysis strategy was applied to the pre-amphetamine data from the DAT:Cilia^-ve^ and their respective wild type littermates. There was a significant effect of time bin on distance traveled (F_(11,396)_=52.09, p<.001), but no main effects or interactions involving sex or genotype (Fs<2.44, ps>.06)(Fig 5A, B). On the stereotypy measure, there was a main effect of time bin (F_(11,396)_=31.36, p<.001) and a time bin x sex interaction (F_(11,396)_=4.52, p=.001), but no main effects or interactions involving genotype (Fs<0.80, ps>.64)(Fig 5C, D). To summarize, neither the GAD2:Cilia^-ve^ nor the DAT:Cilia^-ve^ mice appeared to exhibit abnormalities in baseline activity.

**Figure 4.**
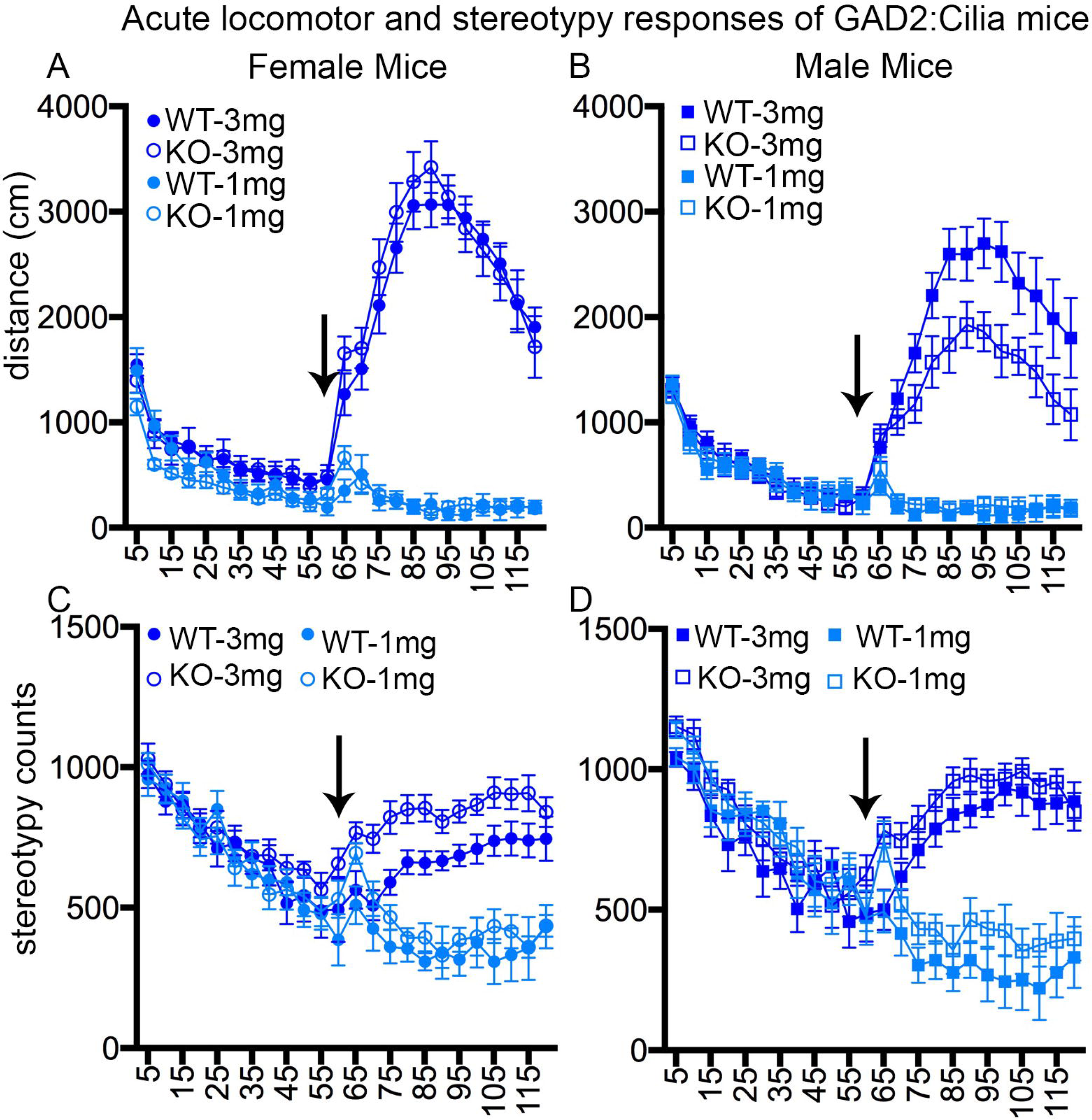
Locomotor and stereotypy responses of GAD2:Cilia mice to acute injections of amphetamine. (A-B) Time course of horizontal activity per 5min bin of wildtype (WT, filled symbol) and GAD2:Cilia^-ve^ (KO, open) mice, before and after 3mg (dark blue) or 1mg/kg (light blue) amphetamine (injection timepoint marked with arrow). Male KO display significantly reduced locomotor activity. (C-D) Time course of stereotypy counts per 5 min bin before and after amphetamine injection (arrow).

**Figure 5.**
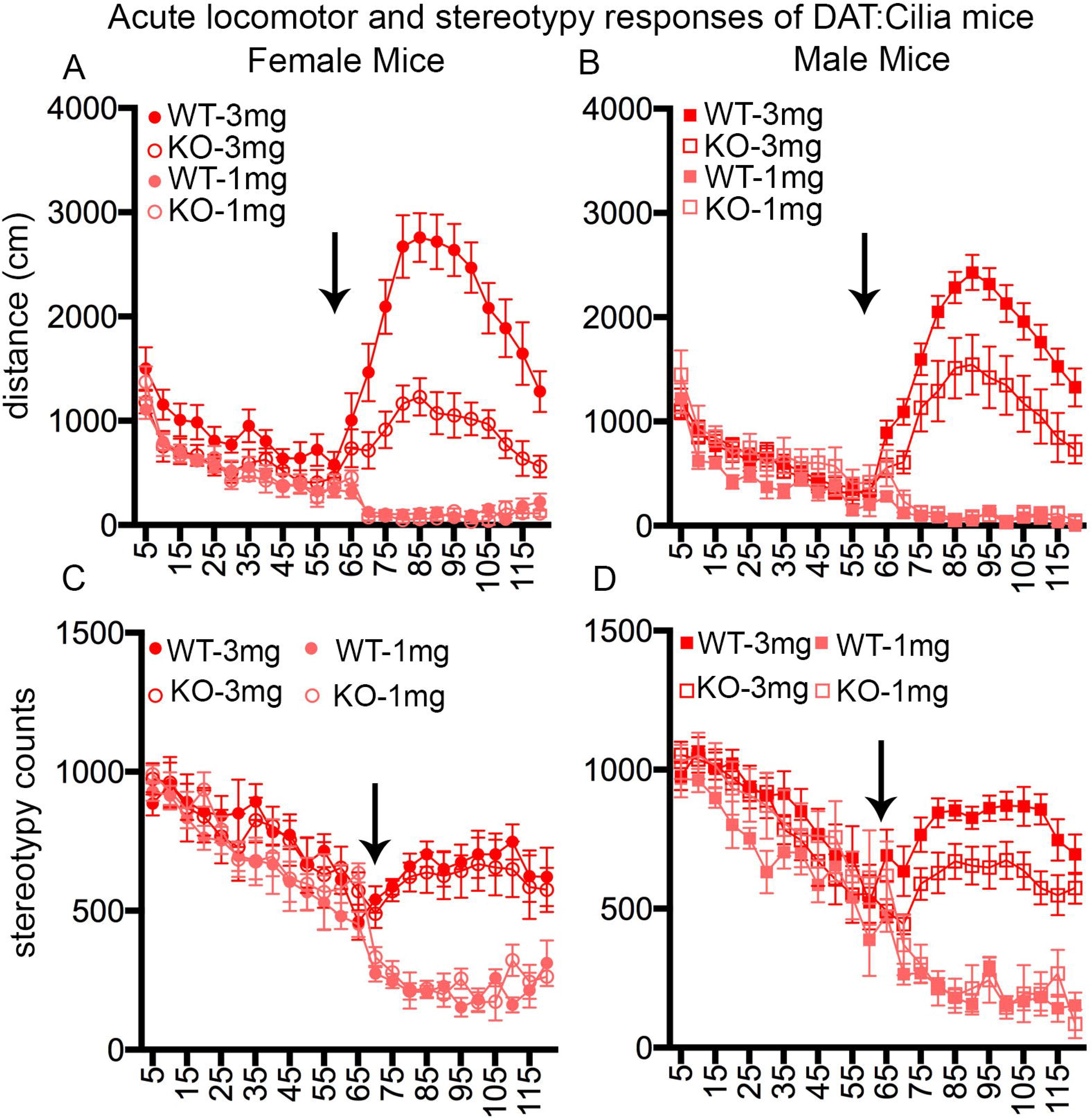
Locomotor and stereotypy responses of DAT:Cilia mice to acute injections of amphetamine. (A-B) Time course of horizontal activity per 5min bin of wildtype (WT, filled symbol) and DAT:Cilia^-ve^ (KO, open) mice, before and after 3mg (dark blue) or 1mg/kg (light blue) amphetamine (injection timepoint marked with arrow). Female (A) and male (B) DAT:Cilia^-ve^ mice show significant reduction in distance traveled following injection of 3mg/kg amphetamine. (C-D) Time course of stereotypy counts per 5 min bin before and after amphetamine injection (arrow).

### 3.3 Effects of acute amphetamine on activity

#### GAD2:Cilia^-ve^ mice

A 3-factor ANOVA (sex x genotype x time bin) conducted on data from the 60 min following 3.0 mg/kg amphetamine administration to GAD2:Cilia^-ve^ mice and their wild type littermates revealed the expected significant effect of time bin (F_(11,418)_=52.51, p<.001), such that amphetamine increased distance traveled, with a peak at approximately 30 min post-injection. Across genotypes, distance traveled was greater in females than males, as indicated by both a main effect of sex (F_(1,38)_=13.16, p=.001) and a sex x time bin interaction (F_(11,418)_=1.86, p=.04)(Fig 4A, B). Most importantly, there was a significant genotype x time bin interaction (F_(11,418)_=2.31, p=.01), such that, across sexes, GAD2:Cilia^-ve^ mice traveled shorter distances than their wild type littermates, particularly in the latter half of the session. Despite the numerically greater effect of genotype in males compared to females, the sex x genotype interaction did not reach statistical significance (F_(1,38)_=3.43, p=.07). Analysis of the stereotypy measure revealed an analogous pattern of results, with a main effect of time bin (F_(11,418)_=23.35, p<.001) and both sex x time bin (F_(11,418)_=2.86, p=.001) and genotype x time bin (F_(11,418)_=1.87, p=.04) interactions (Fig 4C, D). Notably (and in contrast to the distance traveled measure), stereotypy counts were greater in GAD2:Cilia^-ve^ mice compared to wild type, particularly in the first half of the session. Following the first administration of the 1.0 mg/kg dose of amphetamine, there was a significant main effect of time bin (F_(11,308)_=8.92, p<.001) but no other main effects or interactions on the distance traveled measure (Fs<1.39, ps>.17)(Fig 4A,B). There was also a main effect of time bin on the stereotypy measure (F_(11,308)_=7.95, p<.001), as well as a main effect of genotype (F_(1,28)_=4.95, p=.03) such that GAD2:Cilia^-ve^ mice showed greater stereotypy counts than wild type, but no other main effects or interactions (Fs<0.65, ps>.48) (Fig 4C, D).

#### DAT:Cilia^-ve^ mice

A 3-factor ANOVA (sex x genotype x time bin) conducted on data from the 60 min after 3.0 mg/kg amphetamine administration to DAT:Cilia^-ve^ mice and their wild type littermates revealed the expected significant effect of time bin (F_(11,396)_=36.19, p<.001), such that amphetamine increased distance traveled, with a peak at approximately 30 min post-injection. In contrast to the data from GAD2:Cilia^-ve^ mice, there were no sex differences in distance traveled post-amphetamine (main effect of sex F_(1,36)_=0.05, p=.83; sex x time bin interaction, F_(11,396)_=0.88, p=.56). There were, however, robust genotype differences (main effect of genotype, F_(1,36)_=25.25, p<.001; genotype x time bin interaction, F_(11,396)_=4.53, p<.001), such that the locomotor-potentiating effects of amphetamine were attenuated in DAT:Cilia^-ve^ mice (Fig 5A, B). Analysis of the stereotypy measure revealed an analogous pattern of results, with a main effect of time bin (F_(11,396)_=13.06, p<.001) as well as a main effect of genotype (F_(1,36)_=8.89, p=.005), such that stereotypy counts were lower in DAT:Cilia^-ve^ compared to wild type mice (Fig 5C, D). Although this effect was numerically greater in males compared to females, the sex x genotype interaction did not reach statistical significance (F_(1,36)_=3.06, p=.09). Following the first administration of the 1.0 mg/kg dose of amphetamine, there was a significant main effect of time bin (F_(11,231)_=20.18, p<.001), such that distance traveled decreased across the session (Fig 5A, B). There were also interactions between both sex and time bin (F_(11,231)_=1.95, p=.04) and genotype and time bin (F_(11,231)_=2.67, p=.003). The effects of the 1.0 mg/kg dose on the stereotypy measure paralleled those on the distance traveled measure, in that there was also a main effect of time bin (F_(11,231)_=23.93, p<.001), as well as sex x time bin (F_(11,231)_=1.85, p=.047) and genotype x time bin (F_(11,231)_=2.17, p=.02) interactions (Fig 5C, D). In the absence of apparent stimulant effects of amphetamine at this dose, however, it is unclear how to interpret these interactions.

### 3.3 Effects of repeated amphetamine on activity

#### GAD2:Cilia^-ve^ mice

To determine how cilia loss on GAD2-expressing cells affects amphetamine-induced neuroplasticity, activity in response to 5 days of daily amphetamine administration (1.0 mg/kg) was assessed. A 3-factor repeated measures ANOVA (sex x genotype x day) conducted on data summed across the 60 min following each amphetamine injection revealed a main effect of day (F_(4,112)_=41.08, p<.001), such that mice increased their distance traveled across days (i.e., exhibited sensitization to the locomotor stimulant effects of amphetamine, Fig 6A). Although GAD2:Cilia^-ve^ mice exhibited numerically greater distance traveled than their wild type littermates, this effect did not reach statistical significance (F_(1,28)_=2.97, p=.10). No other main effects or interactions were statistically significant (Fs<1.77, ps>.14). The same analysis conducted on the stereotypy measure also revealed a main effect of day (F_(4,112)_=28.68, p<.001), such that stereotypy counts increased across days (Fig 6B). In contrast to the distance traveled measure, however, there were also significant main effects of genotype (F_(1,28)_=14.61, p=.001), such that GAD2:Cilia^-ve^ mice showed greater stereotypy than wild type controls, and sex (F_(1,28)_=14.83, p=.001), such that males showed greater stereotypy than females, as well as a significant sex x day interaction (F_(4,112)_=8.60, p<.001), such that the increase in stereotypy across days was greater in males than females. Considered together, these data suggest that cilia deletion on GAD2-expressing neurons enhances amphetamine-induced plasticity of the locomotor response to the drug.

**Figure 6.**
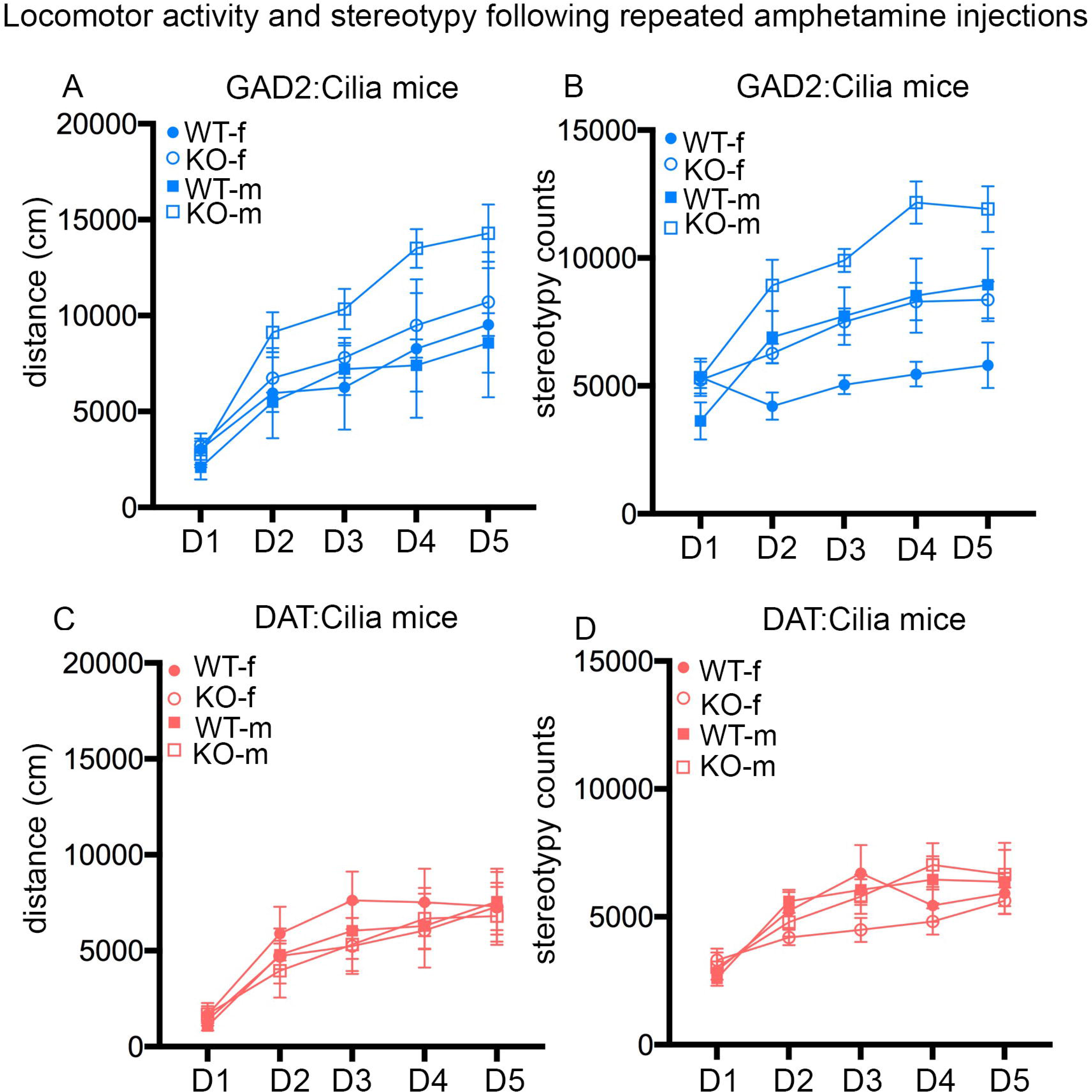
Locomotor and stereotypy responses of GAD2:Cilia and DAT:cilia mice to repeated injections of 1mg/kg amphetamine. Sensitization with 1mg/kg Amp for 5 days. (A-B) Male GAD2:Cilia^-ve^ mice displayed significantly enhanced sensitization in both total locomotor activity (A) and stereotypy counts (B), while female GAD2:Cilia^-ve^ mice show enhancement of just stereotypy counts(B). (C) Total distance traveled in 60min post amphetamine injection each day for DAT:Cilia^-ve^ shows no difference in sensitization of locomotor activity. (D) Total stereotypy counts over 60min post amphetamine injection each day. DAT:Cilia^-ve^ mice show no difference in sensitization of stereotypy counts.

#### DAT:Cilia^-ve^ mice

A 3-factor repeated measures ANOVA (sex x genotype x day) conducted on data from the DAT:Cilia^-ve^ line revealed a main effect of day on distance traveled (F_(4,84)_=28.22, p<.001), such that mice increased their distance traveled across days (Fig 6C). No other main effects or interactions approached statistical significance (Fs<0.56, ps>.57). The same analysis conducted on the stereotypy measure also revealed a main effect of day (F_(4,84)_=29.90, p<.001), such that stereotypy counts increased across days (Fig 6D). The sex x day and genotype x day interactions both approached but did not reach statistical significance (F_(4,84)_=2.10, p=.09 and F_(4,84)_=2.15, p=.08, respectively). No other main effects or interactions approached statistical significance (Fs<0.89, ps>.35). Considered together, these data suggest that cilia deletion on DAT-expressing neurons has minimal if any effects on amphetamine-induced plasticity of the locomotor response to the drug.

## 4. Discussion

Neuronal cilia have been implicated in a variety of developmental processes including dendritic branching and axon targeting [5, 11, 13, 33]. Despite these studies, relatively little is known about the functions of cilia on mature neurons. Given that neurons maintain these structures and enrich them with selective GPCRS, cilia are likely to have important roles in neuronal function. Previous work has identified roles for neuromodulatory GPCRs that are enriched on cilia in mediating behavioral and cellular responses to both stimulants and alcohol, but the specific roles of cilia themselves in these responses has not been studied. Additionally, cilia are dynamic organelles whose structure can be altered through changes in dopamine levels and psychotomimetics [36, 37]. Here we report, to the best of our knowledge, the first behavioral characterization of responses to psychostimulants in animals that lack cilia on specific neuronal populations.

Cilia are critical organelles for development, with numerous cilia-related mutations resulting in embryonic or neonatal lethality [38, 39]. In the current study, targeted ablation of cilia on specific neuron populations post-differentiation did not produce any noticeable changes in survival. A common phenotype among ciliopathies is the development of obesity [34]. We found that removal of cilia from dopaminergic neurons did not alter body weight at 10-12 weeks of age. In contrast, we found that cilia loss on GABAergic neurons resulted in a decrease in body weight. A similar finding was recently reported for mice lacking MCHR1 in GABAergic neurons [40], suggesting that cilia ablation in the present study altered MCHR1 signaling in these neurons and its consequent effects on body weight. Whether the body weight phenotype is due to decreases in food consumption or changes in activity would need to be determined in future studies, though it is notable that the GAD2:Cilia^-ve^ mice exhibited normal activity under baseline conditions.

In both strains, cilia loss did not cause changes in baseline locomotor activity, suggesting that cilia loss in these models does not cause gross abnormalities in motor function. Importantly, however, locomotor activity in the present studies was assessed during the light phase. Previous work has shown that selective loss of MCHR1 on GABAergic neurons does not alter locomotion during the light phase, but does enhance locomotion during the dark phase [40], suggesting that a similar phenotype might be observed in GAD2:Cilia^-ve^ mice. Future studies would be needed to evaluate this possibility, and to determine whether it is evident in dopaminergic cilia knockout mice as well.

The loss of primary cilia on GABAergic and dopaminergic neurons produced both similar and distinct effects on the locomotor responses to amphetamine. DAT:Cilia^-ve^ mice showed marked reductions in locomotor activity (distance traveled) in response to acute administration of a 3.0 mg/kg but not a 1.0 mg/kg dose of amphetamine. This effect was not accompanied by increased stereotypy, suggesting that the decrease in distance traveled was not due to competition from an increase in stereotyped behaviors but instead a reduction in sensitivity/responsivity to the drug. Despite the reduced responses to acute amphetamine, DAT:Cilia^-ve^ mice were no different from wild type controls in their sensitization of responses to repeated amphetamine, suggesting that this form of amphetamine-induced neuroplasticity is not dependent upon dopaminergic neuronal cilia for expression. Like DAT:Cilia^-ve^ mice, GAD2:Cilia^-ve^ mice also showed a reduction in distance traveled in response to acute administration of 3.0 mg/kg but not 1.0 mg/kg amphetamine. The magnitude of this reduction was numerically greater in males than females. Although the interaction with genotype did not reach statistical significance, the data highlight the importance of testing both sexes. Additionally, unlike DAT:Cilia^-ve^ mice, GAD2:Cilia^-ve^ mice showed an increase in stereotypy in response to acute amphetamine. Although it is possible that the reduction in distance traveled was due to an increase in stereotyped behaviors, the fact that the magnitude of the latter effect was greater in females than males suggests that that the genotype-induced changes in distance traveled and stereotypy were not causally linked. Finally, GAD2:Cilia^-ve^ mice showed enhanced sensitization to the stimulant effects of repeated amphetamine. This enhanced sensitization reached statistical significance only for the stereotypy measure and not for the distance traveled measure. GAD2:Cilia^-ve^ mice exhibited a numerical increase in both behaviors, however, suggesting that loss of cilia on GAD2-positive neurons enhances establishment or expression of amphetamine-induced neuroplasticity. This enhanced sensitization to the stimulant effects of amphetamine is similar to that observed in mice in which MCHR1 is constitutively deleted. These mice exhibit enhanced sensitization to some (though not all) doses of amphetamine [41, 42], suggesting that ciliary MCHR1 signaling is involved in the neuroplastic processes supporting sensitization. Importantly, unlike GAD2:Cilia^-ve^ mice, MCHR1 knockout mice also exhibit increased locomotor activity at baseline and in response to acute amphetamine, suggesting that loss of ciliary MCHR1 does not account for the entire MCHR1 knockout phenotype.

There are a number of limitations to the current results. Most obviously, the fact that cilia were ablated on all neurons expressing DAT or GAD2 precludes identification of the specific brain regions in which signaling in this organelle regulates amphetamine responses. Given the central role of midbrain dopaminergic neurons in motor responses to amphetamine, it is likely that the effects observed in DAT:Cilia^-ve^ mice were mediated through this brain region [23]. The widespread distribution of GAD2 neurons, however, renders specifying the anatomical substrates of the effects in GAD2:Cilia^-ve^ mice more challenging. The nucleus accumbens is an obvious potential locus given its important role in the stimulant effects of amphetamine [43]. The stimulant actions of amphetamine are not limited to the nucleus accumbens, however, particularly in the case of psychomotor sensitization, which involves several additional brain systems [44]. Future experiments employing more anatomically-specific approaches (e.g., virally-mediated cilia ablation) will be needed to address this issue. A second caveat to the current data concerns the role of weight loss in the locomotor alterations in GAD2:Cilia^-ve^ mice. Changes in food intake and food restriction can alter responses to both acute and repeated amphetamine [45-48], suggesting that shifts in the responses to amphetamine in GAD2:Cilia^-ve^ mice could have been due in part to their reduced body weight. Although, at least in rats, the effects of food restriction is usually associated with augmentation of locomotor responses. As food restriction is known to increase MCH levels, which act through MCHR1 receptors localized to cilia, these results may provide further support for the necessity of ciliary localization for proper receptor function given a reduction in locomotor responses to amphetamine. Previous work as found that amphetamine induced conditioned place preference is enhanced by food-restriction, but this enhancement is absent in mice lacking MCHR1 [45]. Additionally, the fact that DAT:Cilia^-ve^ mice also exhibited a reduced locomotor response to amphetamine in the absence of body weight differences indicates that a reduction in body weight is not obligate for cilia ablation-induced changes in amphetamine responses.

It is important to note that the effects of cilia ablation on locomotor responses to amphetamine may not necessarily reflect changes in the rewarding or reinforcing properties of the drug. Although manipulations that influence locomotor responses to drugs of abuse can have parallel effects on reward/reinforcement [49], it will be important in future studies to evaluate the effects of cilia ablation in models that more closely align with substance use. Given the demonstrated role of several cilia-enriched GPCRs in substance use [16, 19-21, 41, 50-52], the current data suggest that targeting neuronal cilia could be a useful strategy for reducing drug-seeking behavior.

## Conflict of Interests

The authors report no conflicts of interest.

## Funding

This work was supported by a University of Florida McKnight Brain Institute Pilot Award (JCM), NIH DA047623 (JCM and BS) and NIH T32DC015994 (KRJ).

## Acknowledgments

JCM and BS designed the project. CR, JBR, KRJ, TWT, and JCM performed experiments. JCM and BS analyzed data and wrote the manuscript with input from other authors. We thank Shelby Blaes for assistance with this project, and the NIDA Drug Supply Program for kindly providing d-amphetamine sulfate for these experiments.

